# Comparative study between radiofrequency-induced and muscimol-induced inhibition of cultured networks of cortical neuron

**DOI:** 10.1101/2022.04.05.487108

**Authors:** Clément E. Lemercier, André Garenne, Florence Poulletier de Gannes, Corinne El Khoueiry, Delia Arnaud-Cormos, Philippe Levêque, Isabelle Lagroye, Yann Percher-ancier, Noёlle Lewis

## Abstract

Previous studies have shown that spontaneously active cultured networks of cortical neuron grown planar microelectrode arrays are sensitive to radiofrequency (RF) fields and exhibit an inhibitory response more pronounced as the exposure time and power increase. To better understand the mechanism behind the observed effects, we aimed at identifying similarities and differences between the inhibitory effect of RF fields (continuous wave, 1800 MHz) to the γ-aminobutyric acid type A (GABA_A_) receptor agonist muscimol (MU). Inhibition of the network bursting activity in response to RF exposure became apparent at an SAR level of 28.6 W/kg and co-occurred with an elevation of the culture medium temperature of ~1 °C. Exposure to RF fields preferentially inhibits bursting over spiking activity and exerts fewer constraints on neural network bursting synchrony, differentiating it from a pharmacological inhibition with MU. Network rebound excitation, a phenomenon relying on the intrinsic properties of cortical neurons, was observed following the removal of tonic hyperpolarization after washout of MU but not in response to cessation of RF exposure. This implies that hyperpolarization is not the main driving force mediating the inhibitory effects of RF fields. At the level of single neurons, network inhibition induced by MU and RF fields occurred with reduced action potential (AP) half-width. As changes in AP waveform strongly influence efficacy of synaptic transmission, the narrowing effect on AP seen under RF exposure might contribute to reducing network bursting activity. By pointing only to a partial overlap between the inhibitory hallmarks of these two forms of inhibition, our data suggest that the inhibitory mechanisms of the action of RF fields differ from the ones mediated by the activation of GABA_A_ receptors.

## Introduction

Radiofrequencies are electromagnetic waves ranging from 300 kHz to 300 GHz widely used in modern telecommunication technology. The rapid and continuous increase of environmental man-made RF electromagnetic fields (EMF) has raised concerns about their potential risks on human health. In particular, a large body of research has investigated the possible effects of exposure to RF fields used by mobile phones (300-3000 MHz) on the human central nervous system (CNS) (for reviews see [1–3]). Although evidence exists pointing to an effect of RF fields on brain oscillations [4–7] (reviewed in [8]), evoked potentials [9–10] (but see [11]), and glucose metabolism [12], such changes have not been claimed as having any adverse health effects [13–14]. Interaction between RF fields and biological systems are best understood from a thermal perspective [15–16]. However, compelling evidence suggests that RF fields may also interact with biological systems by producing so-called non-thermal effects (for reviews see [17–19], although see [20–21] for critical reviews), but so far no mechanisms or molecular targets have been identified. Understanding the biological mechanism of non-thermal effects of RF fields on the CNS is not only critical in promoting safety but also holds the promise of useful insights for the development of future biomedical and biotechnological applications.

Early research on various neural preparations reported electrophysiological change in response to RF fields [22–26]. Since then, investigations most frequently indicate that RF fields cause neural activity to decrease [27–35] (but see [24, 36–38]), although the nature of the observed effects might depend on the frequency bands to which the neural preparation is exposed (for example see [28, 38]). In recent years, our laboratory has developed an experimental setup allowing exposing spontaneously active cultures of cortical neurons grown on a planar microelectrode array (MEA) to RF fields, and simultaneously recording the effects [39]. The results obtained with this system indicate that network bursting activity decreases when exposed to RF fields [27] and that the inhibitory response is a function of exposure time and power [28]. Experiments done with equivalent thermal heating suggested that the inhibitory effects of RF fields may originate in part from non-thermal interaction with the nervous tissues. However, the mechanism of action of RF fields on neural networks has remained elusive.

In the present study, we have aimed to contribute to the understanding of the mechanisms of action behind the inhibitory effects of RF fields on cultured cortical neural networks by performing a direct comparison with the inhibitory effects of the GABA_A_ receptor agonist, muscimol (MU). The GABAA receptor is the major inhibitory neurotransmitter receptor responsible for fast inhibition in the mammalian brain [40–41]. Signaling at this receptor is well understood, thus making it a solid reference for comparative studies aiming to infer potential mechanisms of action of particular drugs or treatments. Experiments have been carried out on a new MEA device with improved stability during EMF exposure [42] wherein changes in spiking, bursting activity and action potential (AP) waveform in response to RF fields or MU were analyzed and compared. This comparative approach allowed us to identify similarities and differences between these two forms of inhibition and to employ them as a basis for unravelling a potential mechanism of action of the inhibitory effect of RF fields on cultured neural networks.

## Materials and methods

### Animals

Primary cultures of neocortical neurons were prepared from embryos of gestating Sprague-Dawley rats (Charles River Laboratories, L’Arbresle, France). Experiments involved six gestating rats. All procedures were carried out in compliance with the European Community Council Directive for the Care and Use of laboratory animals (2010/63/EU) and protocols were approved by the Bordeaux Ethics Committee for Animal Experimentation (CEEA - 050).

### Preparation of primary neural culture

Preparation of primary neural cultures was carried out using the methods described in [27–28]. In brief, under anesthetics (5% isoflurane), gestating rats were euthanized by cervical dislocation, embryos (at embryonic day 18) were collected, and their cortices were dissected and treated with a papain-based dissociation system (Worthington Biochemical, Lakewood, CO, USA). Following mechanical dissociation and two steps of centrifugation (the second with an albumin-inhibitor solution), the pellet containing cortical cells (glial cells and neurons) was resuspended in a neurobasal culture medium (NBM) supplemented with 2% B-27, 1% GlutaMAX, and 1% penicillin-streptomycin (Fisher Scientific, Illkirch, France). The recording chips of autoclaved MEAs (Multi Channel Systems MCS GmbH, Reutlingen, Germany) previously coated with polylysine and laminin (Sigma-Aldrich, St. Quentin-Fallavier, France) were plated with a drop of cellular suspension containing 10^5^ cells. Cells were left to sediment and adhere on the MEA chip for up to 2 h and the MEA chambers were then filled with 1 mL of NBM. MEAs were kept in individual petri dishes at 37 °C in a humidified incubator with 5% CO_2_ until mature neural network development. Culture mediums were half-exchanged every 48 h until taking recordings.

### New MEA design and characteristics

In the present study, a modified version [42] of a 60-channel planar MEA introduced in [39] was used. This new design shared the main characteristic of such MEAs, namely the amplifier contact pads placed underneath the printed circuit board, but presented as main evolutions a reduced chip aperture to the limits of the recording zone and several ground planes in the multilayered PCB. These evolutions allowed this device to be steadier in terms of Specific Absorption Rate (SAR) and temperature stability during EMF exposure. Indeed, extensive numerical and experimental dosimetry was carried out to assess SAR values and temperature variation on this new MEA. Although it has been noted that SAR values varied slightly within the culture medium with peak SAR values observed in the vicinity of the electrode tips, microscopic temperature measurements at the electrodes and exposed neurons level did not show any evidence of local temperature hot spots (see [42] for more details on the numerical and experimental dosimetry of the device). In this modified MEA, SAR values normalized per 1 Watt of incident power were estimated at 5.5 ± 2.3 W/kg.

### Electrophysiology and exposure system

The experimental setup for simultaneous electrophysiological recordings and exposure to RF fields or pharmacological agents comprised an MEA coupled to an open transverse electromagnetic cell (TEM) [39, 42–43] and a perfusion system allowing continuous fresh medium exchange with minimal disturbance. RF signal (CW) at 1800 MHz was delivered to the open TEM cell with a signal generator-amplifier (RFPA, Artigues-près-Bordeaux, France). To enable simultaneous recording and exposure to RF fields, MEAs were maintained “sandwiched” between the TEM bottom plate and the preamplifier (MEA1060-Inv, MCS GmbH), as described in earlier publications [27–28, 39, 42]. Once installed on the MEA amplifier, a perfusion holder (MEA-MEM-PL5, ALA Scientific Instruments Inc., Farmingdale, NY, USA) was inserted into the MEA chamber. Perfusion of fresh culture medium was controlled with a peristaltic pump (REGLO ICC, Hugo Sachs Elektronik, March-Hugstetten, Germany) and the optimal perfusion rate (causing minimal disturbance to neural cultures) was set at ~350 μL/min. In these conditions, culture medium was fully exchanged in ~2:50 min. Prior to starting the experiment, cultures were allowed to acclimatize to the continuous medium exchange for ~30 min. Recordings were performed in a dry incubator at 37 °C with 5% CO_2_. Preamplification gain was 1200 and signals were acquired and digitized at 10 kHz/channel with an MCS-dedicated data acquisition board (MC_Card, MCS GmbH). Signals were recorded and visualized with the MC Rack (MCS GmbH) software. After 30 min of baseline recording, neural cultures were exposed for 15 min either to a sham treatment (SH), a pure continuous carrier radiofrequency (RF) at 1800 MHz, or to the GABA_A_ receptor agonist muscimol (MU), (Tocris Bioscience, Bristol, UK). After treatment, post-treatment activity was continuously monitored for 45 min. Data from cultures aged between 17 and 27 days in vitro (DIV) were included in the present study (DIV, Median = 20, Interquartile range, IQR = 4.5, n = 35, all experimental groups collapsed).

### Data analysis and metrics

Processing and analysis of multi-channel data were performed with the software package SPYCODE [44] developed in MATLAB environment (The MathWorks, Inc., Natick, MA, USA). After signal filtering (Butterworth high-pass filter with a cut-off frequency at 70 Hz), spike detection was performed using the differential threshold precision timing spike detection (PTSD) method described by [45] and spike trains were analyzed for burst detection using the method described by [46]. Changes in neural networks activity in response to 15 min of SH, RF or MU exposure were assessed at the level of the entire MEA by pooling data from all active channels (i.e. showing both spiking and bursting activities). Burst detection was used to compute the mean bursting rate (MBR), mean interburst interval (IBI), mean burst duration (BD), mean intraburst spike rate (IBSR), and crossed analysis between burst periods and spike trains allowed computing the mean spiking rate (MSR) for spikes occurring outside bursts. Effects of RF and MU exposure were compared in respect to the SH group after data normalization reflecting the average fractional variation (*R*) of a metric (*M*) during the exposure phase (*M _Expo-sure_*) relative to the baseline reference phase (*M _Baseline_*).

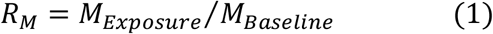

The level of synchronicity for descriptors of bursting activity across MEA channels was evaluated with the coefficient of variation (CV) defined as the ratio (expressed in %) of the average channel standard deviation to the metric mean value (either IBI, BD or IBSR). The lower the CV, the higher synchronization across MEA channels [47–48]. Inter-channel variation for MBR and MSR relative to the overall average fractional variation (i.e. entire MEA) was used to describe the spatial variability of the effects associated with the treatment. This measure was evaluated by computing the normalized root mean square error (*Norm. RMSE*) as follow:

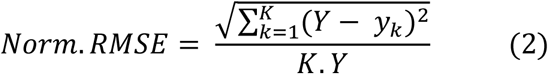

Where ‘*Y*’ is the averaged normalized value of MBR or MSR over all MEA channels (*K*) and ‘*y*’ is the averaged normalized value of MBR or MSR at the level of the individual channel (*k*). For example, a *Norm. RMSE* value equal to 0.5 indicates that the mean inter-channel variation to the mean is of 50 %. Computation methods for the metrics described above are reported in S1 Table.

### AP sorting and waveform analysis

AP detection and sorting were performed with the Offline Sorter V3 (Plexon Inc., Dallas, TX, USA) software over a period of 30 min including 15 min of baseline (pre-exposure phase immediately prior to treatment) and 15 min when neural cultures are continuously exposed to the treatment. To ensure reliable sorting between the two recording phases, pre-exposure and exposure phases were merged into a single file with the MC_dataTool (MCS GmbH) software. Detection threshold was set at five times the standard deviation of the channel noise level and waveform sample-wide containing single event was set at 4 ms (40 sample, 0.8 ms before peak and 3.2 ms after peak). Note that this method of detection differs from the one used in SPYCODE. AP sorting was performed using the T-Dist E-M method (Outlier Threshold 1.5; D.O.F. Mult. 8) and analyses were executed in batch mode. This method enabled detecting on average 67, 991 ± 10,655 (Mean ± SEM) APs per MEA and to sort on average 40,135 ± 6,114

APs per MEA (S1A Fig, data over 15 min during pre-exposure phase from 15 cultures of the RF group used here as representative). Unsorted APs were not analyzed. Hierarchical clustering of the sorted APs indicated that MEA channels presented several sources of AP that were qualified either as major (MAJ), auxiliary (AUX) or minor (MIN) contributors to the total number of sorted spikes (S1A and S1B Figs). On average, MAJ, AUX and MIN AP clusters were respectively observed in 85.3 ± 3.6, 28 ± 4.5, 12.3 ± 1.8 % of the MEA channels and enclosed respectively on average 68 ± 4.4, 21.4 ± 2.5, 10.6 ± 3.3 % of the total amount of sorted APs (S1A Fig). Comparison of the AP timestamps with the burst periods indicated for the MAJ AP cluster that sorted APs inside bursts (_AP_IB) were roughly twice as numerous (~1.9) as sorted APs outside bursts (_AP_OB) and that this proportion decreased to ~1.3 and ~1.1 respectively for the AUX and MIN AP clusters (S1A Fig). As ~89% of the total amount of sorted APs were enclosed in the MAJ and AUX AP clusters, only waveforms from these two clusters were analyzed. The following were measured from these waveforms - peak, anti-peak amplitude, full width at half maximum (FWHM, through linear interpolation), maximum slope of the rising edge and falling edge. Data from MAJ and AUX clusters were then averaged to reflect the overall change in AP waveform in response to the various treatments. Metrics used to quantify changes in AP waveforms are illustrated in S1C Fig and defined in S3 Table.

### Statistics

Statistical analysis was performed using the R software [49] and the ‘PMCMRplus’ library [50]. Unless stated, data in the text and supporting information are reported as median and interquartile ranges (IQR, .i.e. the differences between Q3 and Q1). To evaluate changes relative to the baseline, raw values at baseline for the different metrics showed in Figs 2 and 5 are reported respectively in tabulated form in S2 and S4 Tables. A Kruskal-Wallis test, followed by a Conover’s multiple comparison test, was used to compare differences between groups. A *p*-value < 0.05 was considered statistically significant. Effect size (epsilon-squared, ε^2^), when reported, was calculated with the “rcompanion” [51] R package. Data were plotted with the ‘ggplot2’ [52] and ‘ggpubr’ [53] R packages. The compact letter representation method [54] was used to denote statistical significance after pairwise comparisons with the R package ‘multcompView’ [55]. Pairwise comparisons sharing a common letter are not statistically different but, on the contrary, the ones not sharing any letter are statistically different.

## Results

### Dose response relationship between RF- and MU-induced inhibition

A photograph of the setup illustrating the different parts is shown in Fig 1A. Heating of the culture medium in response to RF exposure at different SAR levels (range: ~4.8 to ~37.9 W/kg) was measured with a fiber optic probe (Luxtron One, Lumasense Technologies, Milpitas, CA, USA; ± uncertainty 0.1 °C) (Fig 1B) immersed in the culture medium under continuous medium exchange (flow rate ~350 μL/min). After 15 min of exposure, heating peaks ranged from ~0.2 to ~1.5 °C respectively for minimum (~4.8 W/kg) and maximum (~37.9 W/kg) tested SAR levels. As cultured networks of cortical neurons are sensitive to RF fields in a dose dependent manner [28], the response relationship between MBR and exposure levels was re-evaluated for the new MEA device used in the present study. With this new type of MEA, inhibition of bursting activity became visible for exposure levels over ~25 W/kg and a reduction of ~50 % in MBR was estimated at ~28.6 W/kg (Fig 1C). At this SAR level, reduction of bursting activity after 15 min of exposure co-occurred with an elevation of the medium temperature of ~1 °C. To compare the effects of RF exposure with those of the GABA_A_ receptor agonist MU under similar levels of inhibition, the relation between MBR and MU concentration was first evaluated (Fig 1D). MU exerts a profound inhibitory action on the activity of cultured cortical networks and its half-maximal inhibitory concentration for the metric MBR (IC_50-MBR_) was estimated to be ~0.25 μM, a value in agreement with other studies on basic receptor and neural culture pharmacology [47, 55–58].

**Fig 1.**
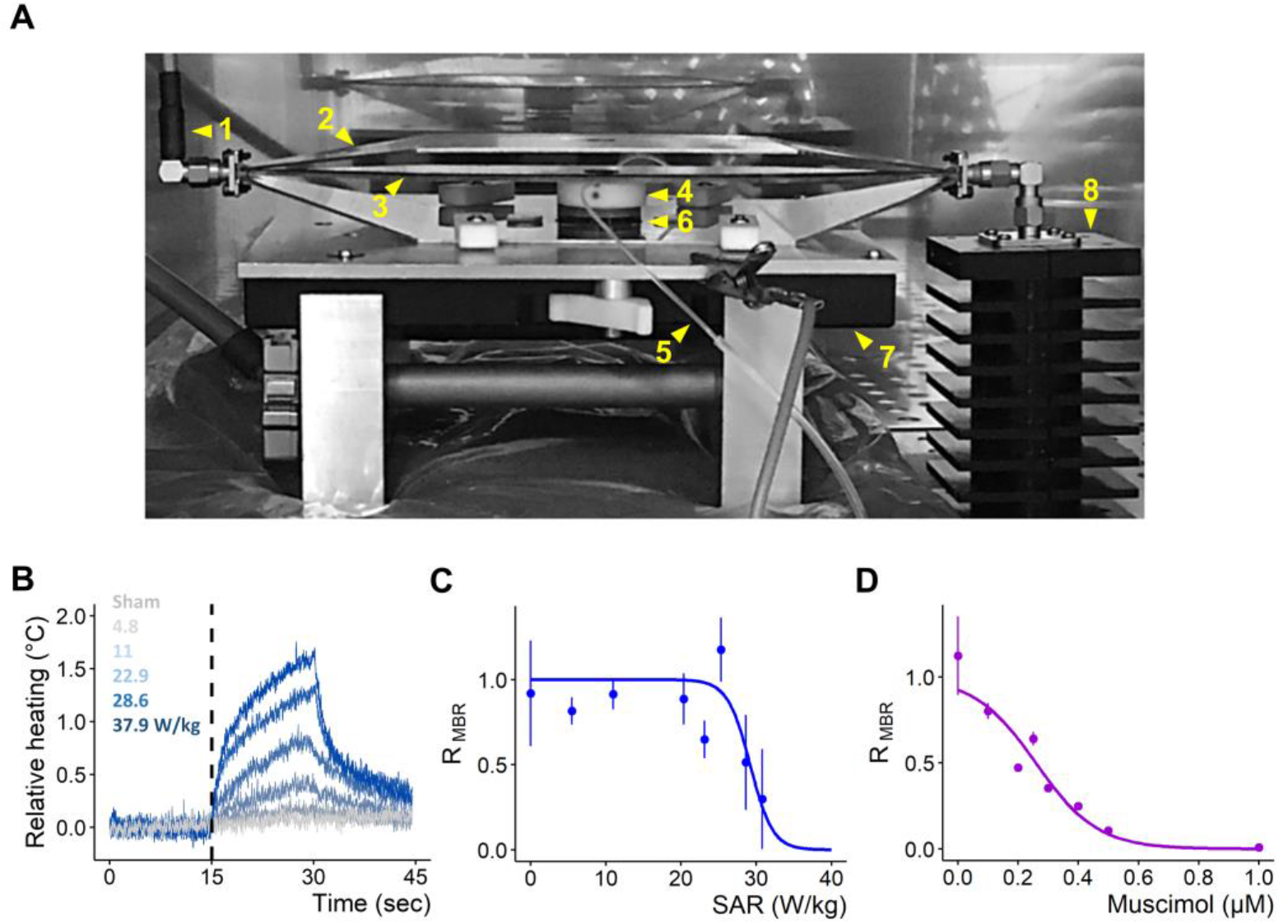
Setup configuration and dose response profile of MBR against SAR level and MU concentration. **(A)** Photograph of the setup configuration used for simultaneous recording on MEA and exposure to RF fields and pharmacological agents. (1) Coaxial cable connecting an RF-generaor/amplifier (located outside the incubator) to (2) an open transverse electromagnetic (TEM) cell. (3) TEM cell septum. (4) Perfusion holder inserted on top of the MEA chamber. (5) Perfusion microtubes for medium exchange. (6) MEA “sandwiched” between TEM bottom ground plate and amplifier ground plate. (7) Inverted MEA preamplifier connected to a MC_Card of a desktop computer. (8) 50 Ω Terminator. **(B)** Relative heating response of the culture medium over 15 min as a function of different SAR levels (W/kg). **(C)** Dose-response relationship between SAR and MBR; results from 21 recordings (18 cultures), 0 (W/kg): n = 21; 5.5: n = 2; 11: n = 3; 20.35: n = 2; 23.1: n = 3; 25.3: n = 3; 28.6: n = 5; 30.6: n = 3. **(D)** Dose-response relationship between MU concentration and MBR; results from 14 recordings (3 cultures), 1e^-4^ (μM): n = 2; 0.1: n = 3; 0.2: n = 1; 0.25: n = 2; 0.3: n = 1; 0.4: n = 1; 0.5: n = 2; 1: n = 2. (C-D) Normalized MBR, ratio of the exposure phase to baseline, data shown as Median ± SD. Fits computed with non-linear least squares method, Pearson’s Goodness-of-Fit: p < 0.05.

**Fig 2.**
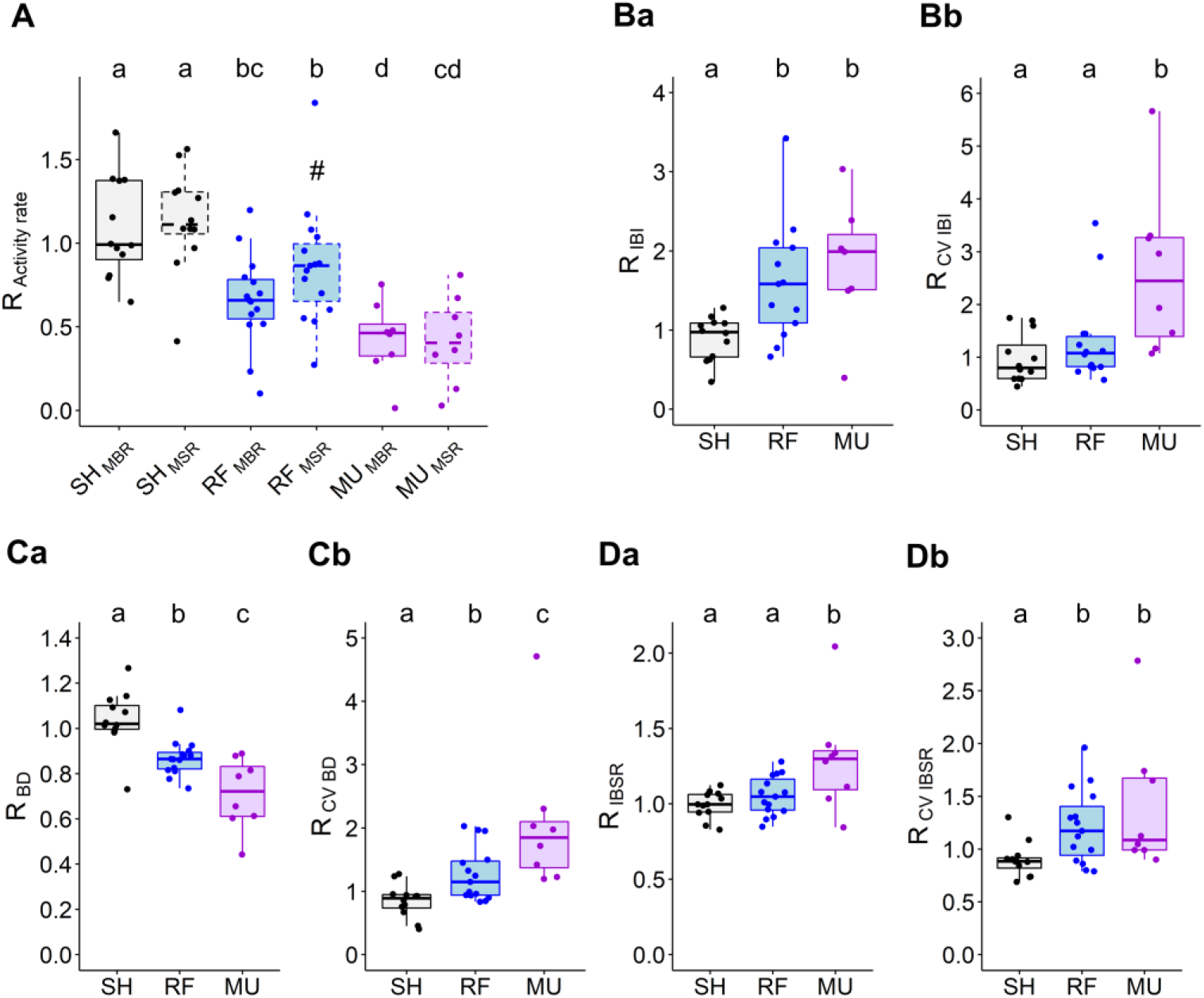
Comparison between RF and MU-induced inhibition of cultured cortical network. **(A)** Average effect of 15 min of exposure to RF and MU on MBR and MSR (spike outside burst periods). Boxplots with dashed box denote MSR data. (#) is indicative of *p* = 0.0535 against RF_-MBR_ and RF_-MSR_. **(Ba)** Average effect of 15 min of exposure to RF and MU on mean inter-burst interval (IBI), **(Ca)** mean burst duration (BD), **(Da)** mean inter-burst spike rate (IBSR). (Bb, Cb and Db) Coefficients of variation (CV) respectively for IBI, BD and IBSR. Normalized data, ratio of the exposure phase to baseline. SH: n = 12; RF: n = 15; MU: n = 8. (A-Db). Lower case letters indicate significant differences between groups.

### RF and MU differentially impacted network activity patterns

The inhibitory effects of RF fields and MU were then compared with respect to an SH group after data normalization (see materials and methods). Definitions of the metrics used to describe changes in network activity in Fig 2 are reported in S1 Table. To assess the magnitude of the reported normalized effects with respect to the raw data, raw data at baseline relative to Fig 2 are tabulated in S2 Table.

Exposure to RF fields (SAR of 28.6 W/kg) or MU (0.25 μM) both reduced MBR (RF: ~35% reduction, SH/RF,*p* < 0.001; MU: ~57% reduction, SH/MU, *p* < 0.001) and MSR (RF: ~14% reduction, SH/RF, *p* < 0.001; MU: ~58% reduction, SH/MU, *p* < 0.001). Inhibitory effects of MU on bursting and spiking activities were on average stronger than for RF exposure (RF_-MBR_/MU_-MBR_, *p*= 0.0412; RF_-MSR_/MU_-MSR_, *p* < 0.001 - Fig 2A). In comparison to MU, RF fields showed a tendency to preferentially inhibit bursting over spiking activity whereas MU reduced equivalently both types of activity (RF_-MBR_/RF_-MSR_, *p* = 0.0543, ε^2^_RF-MBR_ = 0.387, ε^2^_RF-MSR_ = 0.267; MU_-MBR_/ MU_-MSR_, *p* = 0.9057, ε^2^_MU-MBR_ = 0.692, ε^2^_MU-MSR_ = 0.607).

Inhibition of neural network activity was evaluated in the spatial domain by quantifying the inter-channel variability of MBR and MSR variations across all channels of the MEA layout by computing the normalized root mean square error (Norm. RMSE, see materials and methods). Intrinsic variations of this measurement observed in response to SH exposure indicated on average that the level of spatial variability for MBR was slightly lower than for MSR (SH_-MBR_ = 0.22 (0.14); SH_-MSR_ = 0.37 (0.30); *p* = 0.0327). RF- and MU-induced inhibition were both associated with a comparable level of spatial variation of bursting activity across the MEA channels (RF = 0.20 (0.22); MU = 0.28 (0.12); *p* = 0.3059). The degree of spatial variability in MBR was not different from the intrinsic spatial variability observed in response to SH exposure (*p* = 0.3024). In the same way as for the data for MBR, the data for MSR indicated that RF- and MU-induced inhibition caused spiking activity to vary equivalently in space (RF = 0.54 (0.16); MU = 0.65 (0.21); *p* = 0.1848) but spatial fluctuations of MSR were higher than for the intrinsic variation observed with SH exposure (*p* = 0.0037, pooled MSR data across RF and MU); although as in SH exposure, spatial variations of MBR were lower than for MSR (*p* < 0.001). Collectively these data indicate that RF-induced inhibition occurred within the MEA space as diffusely as the pharmacological inhibition induced by MU.

Comparison between RF- and MU-induced inhibitions was pursued with descriptors of bursting activity such as IBI, BD and IBSR and their respective indicators of synchronization across MEA channels with the coefficient of variation (CV, see materials and methods and metrics definition in S1 Table). In response to RF and MU, bursting activity becomes increasingly sparse, as seen by increased IBI (SH/RF, *p* = 0.0020, SH/MU, *p* < 0.001; RF/MU, *p* = 0.4528 - Fig 2Ba). Compared to RF exposure, the inhibitory action of MU was accompanied by a desynchronization of bursting activity across MEA channels as seen by an increased CV IBI (SH/RF, *p* = 0.2318; SH/MU, *p* < 0.001; RF/MU, *p* = 0.0098 - Fig 2Bb). RF and MU both decreased BD (SH/RF, *p* < 0.001; SH/MU, *p* < 0.001; RF/MU, *p* = 0.0178 - Fig 2Ca) and desynchronized BD across MEA channels (CV BD: SH/RF, *p* = 0.0035, SH/MU, *p* < 0.001 - Fig 2Cb) but this effect was of a higher magnitude for MU (RF/MU, *p* = 0.0128). MU, but not RF exposure, increased IBSR (SH/RF, *p* = 0.2919; SH/MU, *p* = 0.0069; RF/MU, *p* = 0.0476 - Fig 2Da). However, both treatments desynchronized IBSR across MEA’s channels (CV IBSR: SH/RF, *p* = 0.0062; SH/MU, *p* = 0.0042; RF/MU, *p* = 0.5388 - Fig 2Db).

### Differential effect of RF and MU on neural networks temporal activity pattern

Analysis and comparison of the two forms of inhibition were pursued in the temporal domain by measuring bursting rate (BR) and spiking rate (SR) over time (Fig 3). In response to RF or MU, BR dramatically decreased by about half of the baseline level within the first minute following exposure (Fig 3A). Similarly to BR, SR reduced within the first minute following exposure onset but, in contrast to MU, the latter appeared on average to be less affected by RF fields (Fig 3B). Quantification of the rate of BR inhibition during the initial phase of exposure (initial inhibitory rate, see metrics definition in S1 Table) indicated that RF fields and MU both impacted BR with an equivalent initial potency (SH/RF, *p* < 0.001; SH/MU, *p* = 0.0035; RF/MU, *p* = 0.9243 - Fig 3C). The initial inhibitory rate for SR in response to RF exposure showed a greater level of variability than for BR and was no different from SH (SH/RF, *p* = 0.1741 - Fig 3C). On the other hand, MU inhibited BR and SR with an equivalent initial potency (SH_-SR_/MU_-SR_, *p* < 0.001; MU_-BR_/MU_-SR_, *p* = 0.6850 - Fig 3C). Following the initial action of the treatments, BR and SR showed a tendency for a slight regain of activity, although this effect was more marked for MU. In response to washout of MU, a dramatic short-lasting regain of activity of about 1 min was observed. This phenomenon qualified as a postinhibitory rebound (PIR, see metrics definition in S1 Table) was, on average, visible both for BR and SR (Figs 3A and 3B) but only significantly detected for bursting activity (SH_-PIR-BR_/ MU_-PIR-BR_, *p* = 0.0128; SH_-PIR-SR_ / MU_-PIR-SR_, *p* = 0.0549 - Fig 3D). Interestingly, PIR was not observed in response to RF exposure cessation (SH_-PIR-BR_ / RF_-PIR-BR_, *p* = 0.8420; SH_-PIR-SR_ / RF_-PIR-SR_, *p* = 0.9821 - Fig 3D). Successive recording phases indicated that neuronal network activity fully recovered from treatment and temporally evolved similarly to SH.

**Fig 3.**
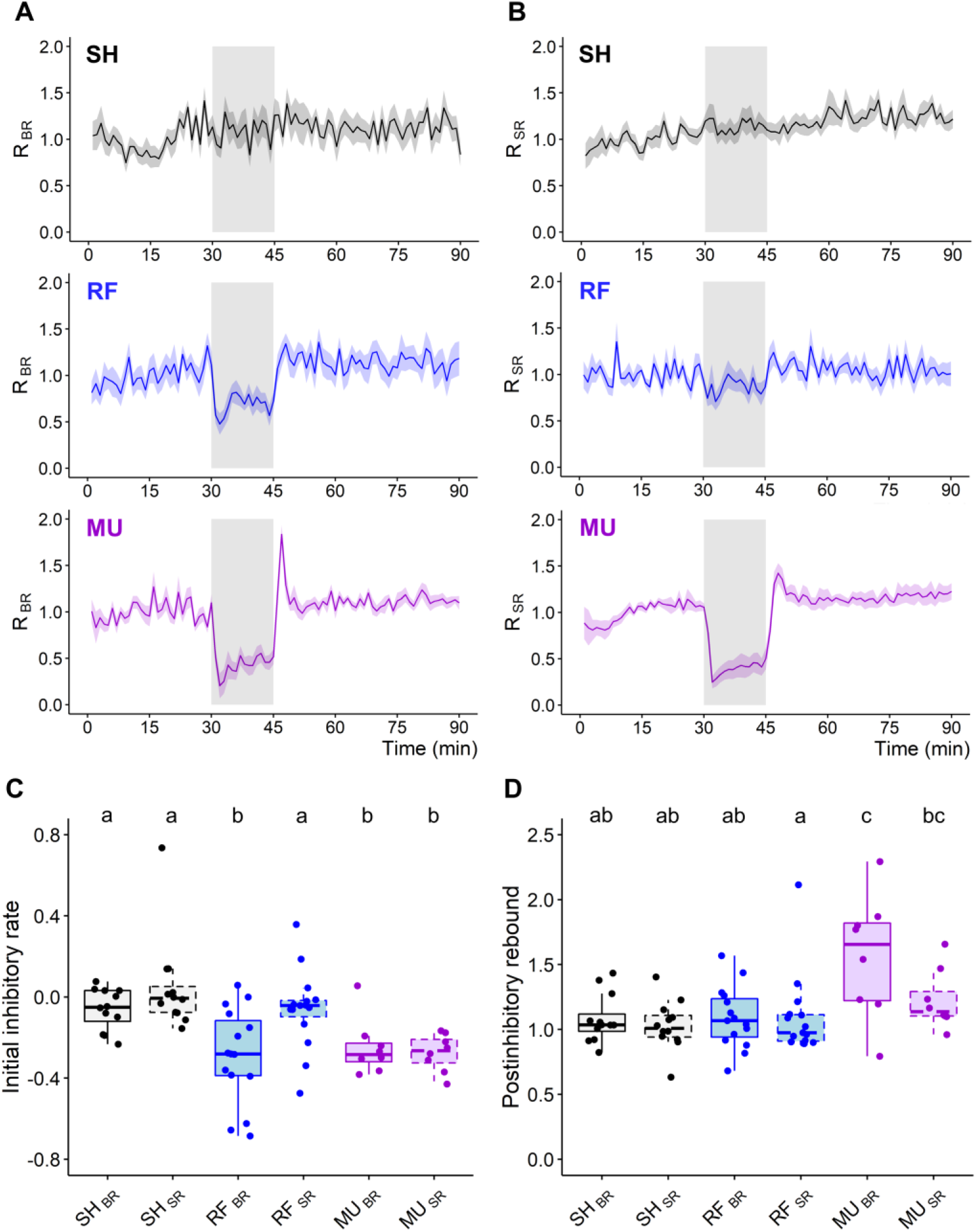
Temporal dynamic of RF and MU-induced inhibition on bursting and spiking rates. **(A-B)** Normalized temporal time course of bursting rate (BR, left) and spiking rate (SR, right) over 90 min for SH (top), RF (middle) and MU (bottom) groups (1 min bin-size, data show as Mean ± SEM). The exposure phase is symbolized by a gray shadowed area. **(C)** Initial inhibitory rate in response to RF and MU exposure. **(D)** Quantification of the postinhibitory rebound in response to treatment cessation. Boxplots with dashed box denote SR data. SH, n = 12; RF, n = 15; MU, n = 8. (C-D). Lower case letters indicate significant differences between groups.

Similarities and differences in the temporal domain between the two treatments are once again exemplified in Figs 4A and 4B with data from two representative cultures exposed either to RF fields or MU. In these examples, the MU experiment is initially marked by an abrupt shutdown of neural activity, lasting a few minutes, followed by a slight and gradual return of activity. Following washout of MU, network BR undergoes a short period of rebound excitation which then re-stabilizes (note the absence of rebound excitation for SR). On the contrary, the RF exposure experiment did not display such dynamics but was rather associated with a strict slow-down of network activity with bursts peaking less frequently above the normalization line.

**Fig 4.**
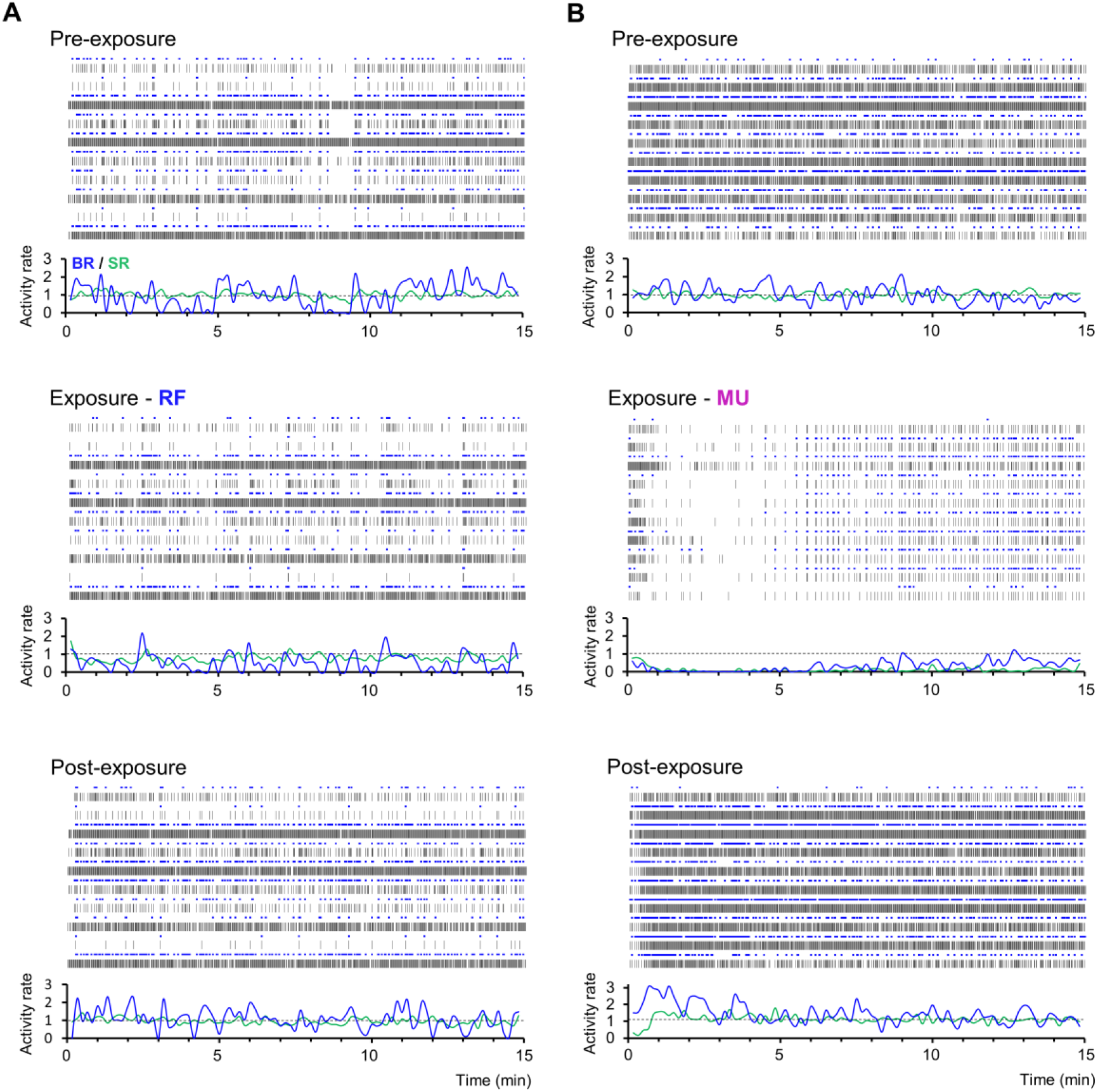
Representative recordings showing the temporal time course of RF- and MU-induced inhibition of neural networks. **(A-B)** Data from 10-selected electrodes of 2 independent cultures either exposed to RF (left) or MU (right) showing spiking (SR) and bursting rate (BR) along three recording segments of 15 min during pre-exposure (top), exposure (middle), and post-exposure recording phases (bottom). Neural activity is shown as spike raster plot capped in blue for markers of burst detection. Below each raster plot is the corresponding normalized BR (blue) and SR (green) computed overtime along non-overlapping sliding windows of 10 sec, dashed lines representing the normalization level.

**Fig 5.**
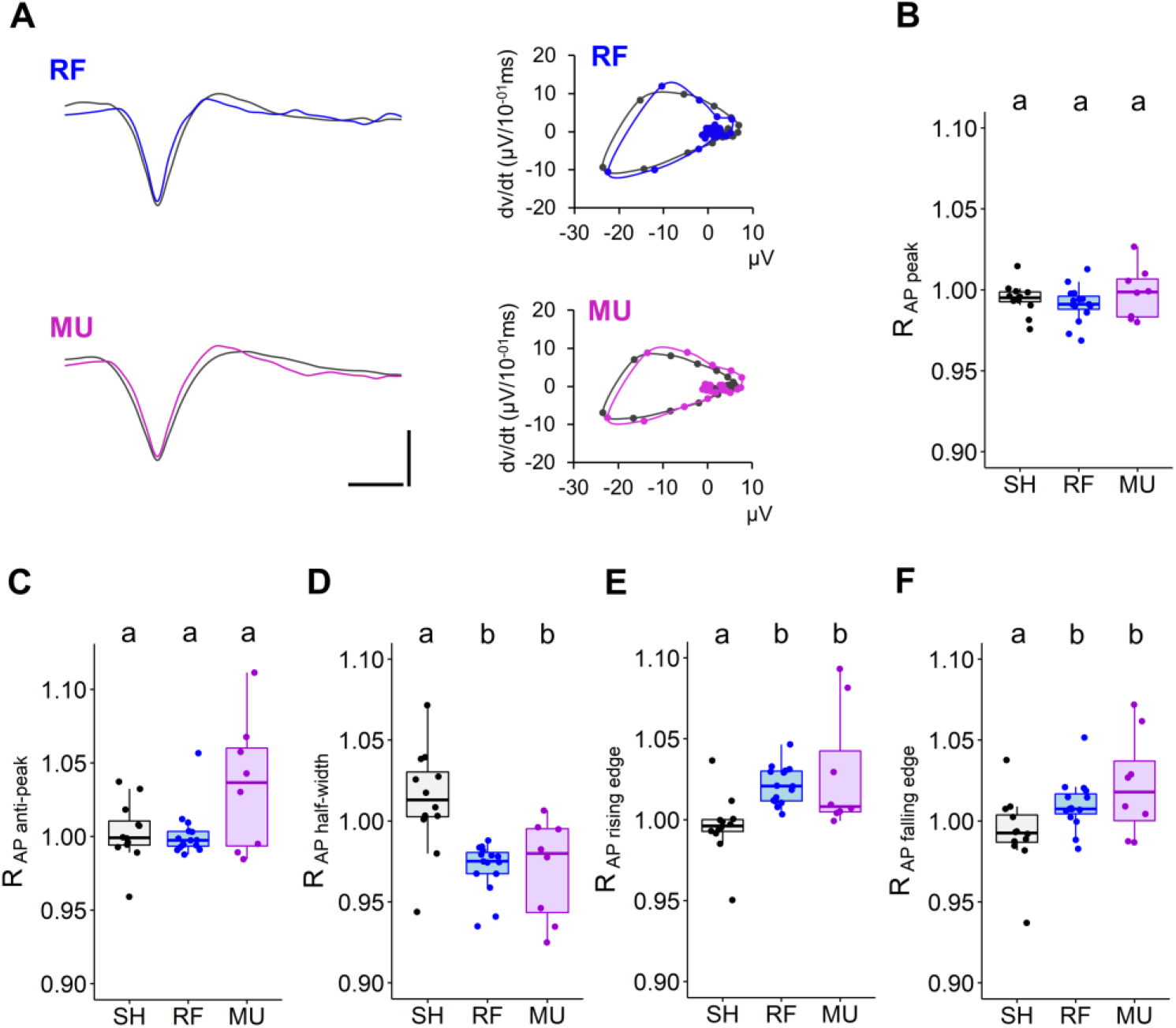
Change in AP waveform in response to RF and MU exposure. **(A)** Representative average AP traces from a single unit (left) and associated phase plot (right) before and during exposure to RF (top) and MU (bottom). Scale: (y): 15 μV; (x): 500 μs. **(B-F)** Boxplots showing variation in AP peak (B) and anti-peak amplitude (C) half-width (D) maximal rising (E) and falling edge (F). Normalized data, ratio of the exposure phase to baseline. SH, n = 12; RF, n = 15; MU, n = 8. (B-F). Lower case letters indicate significant differences between groups.

### RF and MU produce similar AP waveform alteration

The inhibitory effects of RF fields and MU were next analyzed and compared at the level of single-unit activity by evaluating changes in AP waveforms (Fig 5). Definitions of the metrics used to describe changes in AP waveform are illustrated in S1C Fig and defined in S3 Table. To assess the magnitude of the reported normalized effects in respect to the raw data, raw data at baseline relative to Fig 5 are tabulated in S4 Table. After hierarchical clustering of spike events, data from the two main AP clusters were analyzed in a pooled manner (see materials and methods section and S1 Fig for more details on AP detection, sorting, cluster repartition and waveform analysis).

The average effects on AP waveform in response to RF fields and MU are shown from two representative single units and their respective phase plots in Fig 5A. Analysis of AP waveforms showed that, in respect to SH, RF and MU exposure neither impacted the AP peak amplitude (*p* = 0.3511 - Fig 5B) nor the anti-peak amplitude (*p* = 0.2859 - Fig 5C), but that both treatments narrowed the AP half-width (SH/RF, *p* < 0.001; SH/MU, *p* = 0.0018; RF/MU, *p* = 0.6065 - Fig 5D). This narrowing effect occurred symmetrically with both depolarization and repolarization phases occurring at a faster rate (slope of the rising edge: SH/RF, *p* < 0.001; SH/MU, *p* = 0.0038; RF/MU, *p* = 0.3547 - Fig 5E; slope of the falling edge: SH/RF, *p* = 0.0374; SH/MU, *p* = 0.0224; RF/MU, *p* = 0.5659 - Fig 5F). As confirmation, phase plots generally show steeper slopes along the AP cycle, albeit of small amplitude. Analysis of the size effect indicated a stronger effect on the rising than on the falling edge of the AP (RF: ε^2^_rising_ = 0.475; ε^2^_falling_ = 0.194; MU: ε^2^_rising_ = 0.384; ε^2^_falling_ = 0.180) suggesting that narrowing of the AP half-width in response to RF and MU exposure occurred primarily through a mechanism that increases the depolarization slope.

## Discussion

In the present study, exposure to RF fields were performed at an SAR level of 28.6 W/kg, a value ~1.4 times lower than levels used in [27–28]. Indeed, a recent re-evaluation of the dosimetry [42] indicated estimated SAR values per Watt of incident power of 5.5 ± 2.3 W/kg and 40.3 ± 5.3 W/kg respectively for the present and earlier MEA versions [27–28]. This re-evaluation was made possible thanks to the continuous progress in experimental and numerical dosimetry and better assessment of influencing environmental factors [42]. The SAR level of 28.6 W/kg is however higher than local basic safety restrictions fixed at 2.0 W/kg [13]. Therefore this study is rather limited regarding the potential adverse effects of man-made environmental RF fields on human health. RF exposure for 15 min at an SAR level of 28.6 W/kg decreased reversibly bursting activity of ~35 % and co-occurred with an elevation of the culture medium temperature of ~1 °C. The activity rate of neural culture is influenced by temperature with hypo- and hyperthermia being respectively associated with lower and heightened neural activity [28, 36, 59–60] but see [61]. In line with data reported in these studies, previous experiments from our lab showed that heating of the culture medium by ~1 °C slightly increased bursting activity [28] thus suggesting that the observed effect of RF fields might have, in part, non-thermal origins.

We have previously reported that exposure to RF fields decreases the bursting activity of cultured networks of cortical neurons [27] and that this inhibitory effect increases as exposure time and SAR levels increase [28]. In the present study, investigations of the inhibitory effects of RF fields were pursued by performing a direct comparison with the effects of the GABA_A_ receptor agonist MU. Our results showed that in contrast to MU, RF exposure preferentially inhibits bursting over spiking activity. Although spiking activity was reduced by RF exposure, inhibition was more variable and weaker than for bursting activity. Other studies with cultured networks of cortical neurons also reported that MU equivalently inhibits spiking and bursting activity [47, 57]. GABAergic inhibition in the brain can be classified as either phasic or tonic [62]. The first depends on fast activation of synaptic GABA_A_ receptors from synaptically released GABA, whereas the second depends on sustained activation of peri- and extrasynaptic GABA_A_ receptors by ambient GABA. In our experiments, continuous application of MU in the culture medium activates both synaptic and peri-extrasynaptic GABA_A_ receptors, which ultimately leads to a tonic neural hyperpolarization. Neuronal excitability is in essence equivalently reduced throughout the network subcomponents and an equivalent reduction in activity patterns based on regular spiking, intrinsically bursting neurons as well on network collective bursting behavior is observed. As RF exposure differentially impacted spiking and bursting activity, one may argue that cell hyperpolarization is not the main force driving the inhibitory effects of RF on neural networks. Studies on the effect of RF exposure on the membrane potential of excitable cells (cardiomyocytes and neurons) has led to conflicting results, with some showing no effect [23, 26, 63–64], others showing hyperpolarization [31], and sometimes both, depending on the region studied after acute exposure of the whole animal [34]. Detailed electrophysiological investigations in our experimental conditions are needed to shed light on this point.

At the cellular level, cortical neurons can generate bursts based on intrinsic properties such as hyperpolarization-activated current (*I*_h_), subthreshold membrane oscillations and T-type calcium current, above which high frequency action potentials fire for a brief period [65–67]. At a network level, bursts can be generated intermittently in a collective manner as an emergent property [68–69] relying on the development of an excitatory-inhibitory oscillating network [70–71]. On that note, possible hypotheses could be that reduced bursting activity in response to RF exposure is due to a predominant action on intrinsically bursting neurons over regular spiking neurons or, alternatively, that the effect of RF manifests itself on a larger scale by reducing network collective bursting behavior. Interestingly, some authors have suggested that the extremely low-frequency EMFs (high-intensity power frequency, 50 Hz) enhance the activity of cultured networks of cortical neurons by modulating the activity of pacemaker-like interneurons [38]. To our knowledge, this research avenue has not yet been further investigated by other laboratories. Nevertheless, our experiments focused on mature neocortical cultures where network bursts substantially contribute to the overall burst count (~60 to ~80% of the total number of bursts) and no discrimination in our analysis was considered between isolated bursts and network bursts. Therefore, the observed inhibition of bursting activity in response to RF exposure mostly originates from a reduction of network collective bursting behavior. RF exposure at different levels of culture maturity (i.e. irregular and slightly synchronized bursting vs. regular and highly synchronized bursting) is of interest to determine whether neural network topology is a factor determining the sensitivity to RF fields. Moreover, detailed analysis with improved detection algorithms could help to better differentiate between the effect of RF exposure on the different network subcomponents and related activity patterns.

Descriptors of neural networks bursting activity were similarly impacted by RF and MU exposure. In the two forms of inhibition, decreased MBR was accompanied by increased IBI and decreased BD, but data suggested that only inhibition induced by MU was accompanied by increased IBSR. However, the reported effect of MU on IBSR seems to contradict the results of a recent thorough study done under similar experimental conditions [47], thus making it difficult to evaluate the pertinence of this observation in comparison to RF exposure. At neural networks level, a shift in the balance between excitation and inhibition strongly contributes to control burst phase, termination and intraburst spiking rate [47–48, 72–73]. Both Inhibition and disinhibition cause a shortening of the BD. The former occurs with reduced IBSR whereas the second occurs with increased IBSR. Indicators of network bursting synchronization were differently impacted by RF and MU exposure. During the two forms of inhibition BD and IBSR synchronization decreased over the network but only MU shifted network bursting behavior from regular and synchronized to more irregular and less synchronized. This observation suggests that the effects of RF exposure exert fewer constraints on network functioning than those mediated by the activation of the GABA_A_ receptor. The desynchronizing effect of MU on network bursting behavior can most likely be attributed to its hyperpolarizing action. Indeed, it has been shown that inverting the polarity of the GABA action, i.e. depolarizing toward hyperpolarizing, can evoke desynchronized premature-like network activity in young, moderately synchronized, cultures [48].

Upon recovery from the inhibitory effects of MU but not from those of RF exposure, networks showed a dramatic regain in bursting activity that persisted recurrently in a synchronous manner for ~1 min. This phenomenon relies most likely on the intrinsic property of cortical neurons’ so-called postinhibitory rebound and refers to the ability of a neuron to generate rebound excitation upon termination of an inhibitory signal [74–75]. Postinhibitory rebound is involved in a variety of basic brain processes such as rhythmic recurrent activity [76] and short-term plasticity [77]. This phenomenon relies on several mechanisms occurring in response to hyperpolarization such as activation of hyperpolarization-activated cyclic nucleotide-gated (HCN) channels and deinactivation of low voltage-activated T-type calcium channels and persistent sodium channels [78–81]. In our conditions, postinhibitory rebound occurred in response to washout of MU and consecutive removal of tonic hyperpolarization. The absence of postinhibitory rebound in response to RF exposure cessation might furthermore imply that RF fields exert their inhibitory effects without hyperpolarizing neurons. Reduced bursting activity combined with the lack of postinhibitory rebound might suggest that RF fields potentially interfere with the functioning of ion channels involved in these modalities such as of HCN, T-type calcium channels and persistent sodium channels. Interestingly, it has been reported that exposure to extremely low-frequency-EMF (50 Hz, 0.2 mT, 1 hour) inhibited T-type calcium channels in mouse cortical neurons [82]. However, no comparison with other types of currents was made, making it difficult to assess the relevance of this observation in the present study (see [83–84] for reviews on EMF and calcium). Nevertheless, the rapid onset of the effects of RF fields and their reversibility are in favor of a mechanism interacting with fast operating targets at the membrane level such as ion channels. For a detailed review on EMF with cell membranes, organelles and bio-molecules see [19]. Thorough investigations with co-exposure of RF fields and pharmacological agents will enable directly testing potential interactions with ion channels.

Analysis of AP waveform showed that RF- and MU-induced inhibition co-occurred with a slight symmetrical narrowing effect of the AP half-width. Although other studies have reported on the narrowing effect of RF exposure on AP waveform [29, 31, 34] (but see [26, 30]), the mechanism of action through which RF fields alter the AP waveform remains to be established.

Changes in the AP half-width exert direct influences on the efficacy of synaptic transmission [85–88] and might contribute to the inhibitory effect of RF exposure on network bursting activity. Commonalities in the changes in AP waveforms in response to RF and MU exposure suggest a potential overlapping mechanism between these two modalities. A possible point of convergence could be a similar effect on the membrane resistance. Indeed, a decrease in membrane resistance in response to MU [89–90] has also been observed in response to RF fields [22–23] but see [25–26, 91] and millimeter waves (MMWs, 30-300 GHz) [29]. The AP shape strongly relates to membrane resistance, with decreased and increased resistance being respectively associated with narrower and broader AP [92–93]. Membrane resistance and AP waveform are also very sensitive to changes in temperature with increased and decreased temperature leading respectively to lower/narrower and higher/broader membrane resistance and AP [92–95]. Therefore, it cannot be excluded that the observed effect on AP waveform has a thermal origin [31]. Recently, it has been reported that mid-infrared radiations also shorten AP by accelerating its repolarization, through an increase in voltage-gated potassium currents [95]. Mechanisms of RF field effects might differ from mid-infrared radiation as they manifest predominantly by a steeper depolarization phase. Detailed electrophysiological experiments combined with accurate temperature control or bulk heating are required to elucidate the mechanism of RF fields on AP waveform. Moreover, the hypothesis that decreased AP half-width contributes to decreased network bursting behavior should be investigated in silico with neural simulation.

## Acknowledgments

The authors thank Stephano Buccelli for his contribution in implementing new scripts in the software package SPYCODE and Prof. Dr. Patrik Krieger for sharing tools for spike sorting.

## Author Contributions

**Conceptualization:** André Garenne, Isabelle Lagroye, Noёlle Lewis.

**Data Curation:** Clément E. Lemercier.

**Formal Analysis:** Clément E. Lemercier.

**Funding Acquisition:** André Garenne, Delia Arnaud-Cormos, Philippe Levêque, Isabelle Lagroye, Noёlle Lewis.

**Investigation:** Clément E. Lemercier.

**Methodology:** Clément E. Lemercier, André Garenne, Florence Poulletier de Gannes, Corinne El Khoueiry, Delia Arnaud-Cormos, Philippe Levêque, Isabelle Lagroye, Yann Percherancier, Noёlle Lewis.

**Project Administration:** Noëlle Lewis.

**Resources:** Noëlle Lewis.

**Supervision:** André Garenne, Noëlle Lewis.

**Visualization:** Clément E. Lemercier.

**Writing – Original Draft Preparation:** Clément E. Lemercier

**Writing – Review & Editing:** Clément E. Lemercier, André Garenne, Florence Poulletier de Gannes, Corinne El Khoueiry, Delia Arnaud-Cormos, Philippe Levêque, Isabelle Lagroye, Yann Percherancier, Noëlle Lewis.

## Data Availability Statement

All relevant data are within the manuscript and its Supporting Information files.

## Supporting information

**S1 Table.** Metrics definition: Analysis of neural network activity

| Analysis of neural network activity |  |
| --- | --- |
| Metrics |  |
| Day in vitro (DIV) | Age of the culture (in days) from preparation date (DIV<sub>0</sub>) to recording date (DIV<sub>n</sub>) |
| Active channel (k) | Number of channels showing both spiking and bursting activities |
| Mean spike rate (MSR) | Sum of channel mean spike rate (sec<sup>-1</sup>) |
| Mean burst rate (MBR) | Sum of channel mean burst rate (sec<sup>-1</sup>) |
| Network bursting rate (NtBR) | Rate of bursts occurring simultaneously in ≥ 20 channels (min<sup>-1</sup>) |
| % of spike outside bursts | Ratio of the number of spikes outside bursts to the total of number of spikes. |
| Mean Interburst interval (IBI) | Pooled mean of mean channel IBI (sec) |
| Mean Burst duration (BD) | Pooled mean of mean channel BD (msec) |
| Mean intraburst spike rate (IBSR) | Pooled mean of mean channel IBSR (n spike in burst / burst duration)*1000), Hz) |
| Coefficient of variation (CV) | Ratio (expressed in %) of the average channels standard deviation to the metric mean value (either IBI, BD or IBSR) |
| Initial inhibitory rate | Linear regression of 4 points over peri-exposure period, 2 min before exposure-onset and 2 min after exposure-onset |
| Postinhibitory rebound | Ratio between the maximal values (either MBR or MSR) retrieved in two consecutive non-overlapping windows of 4 min after exposure-offset |

**S2 Table.** Raw values at baseline of the various metrics used to describe neural networks activity across the various experimental groups.

| Metrics | SH | RF | MU |
| --- | --- | --- | --- |
| N culture | 12 | 15 | 8 |
| N culture per animal | 2 (3) | 4 (3.5) | 3 (1.5) |
| Day In Vitro | 18.5 (1.5) | 20 (2) | 23 (2.5) <sup>*(2)</sup> |
| Active k | 49 (13) | 54 (6) | 54 (5) |
| MSR (sec<sup>-1</sup>) | 15.6 (18.5) | 23.4 (37.3) | 52 (43.2) <sup>*(2)</sup> |
| MBR (sec<sup>-1</sup>) | 2.5 (2.6) | 3.5 (2) | 11.3 (12.7) <sup>*(2)</sup> |
| NtBR (min<sup>-1</sup>) | 3.9 (5.3) | 3.3 (1.6) | 16.9 (16.4) <sup>*(2)</sup> |
| % of spike outside bursts | 28.3 (21.9) | 25 (21.5) | 16.9 (12.3) |
| IBI (sec) | 40.3 (35.6) | 23.9 (11.8) | 6.8 (10.5) <sup>*(2)</sup> |
| CV IBI (%) | 19.3 (13.7) | 17.4 (8.8) | 8.8 (7.7) <sup>ns</sup> |
| BD (ms) | 89 (102.5) <sup>*(1)</sup> | 217 (91.8) | 200.6 (115.4) |
| CV BD (%) | 7 (3.4) | 5.2 (3) | 4.2 (1.8) <sup>*(2)</sup> |
| IBSR (Hz) | 160.3 (46.9) | 131.4 (42) | 122 (35.9) |
| CV IBSR (%) | 6.8 (3.9) | 5.1 (2.1) | 4.2 (2.3) |

Data expressed as Median (IQR). <sup>\*(1)</sup> Indicates significant difference between SH-RF and SH-MU pairs ( $p < 0.05$ ) and <sup>\*(2)</sup> indicates significant difference between MU-SH and MU-RF pairs ( $p < 0.05$ ), <sup>ns</sup> indicates no significant differences between groups. Pairwise comparison done with Kruskal-Wallis test followed by Conover's all-pairs posthoc test. SH, n = 12; RF, n = 15; MU, n = 8.

**S1 Fig.**
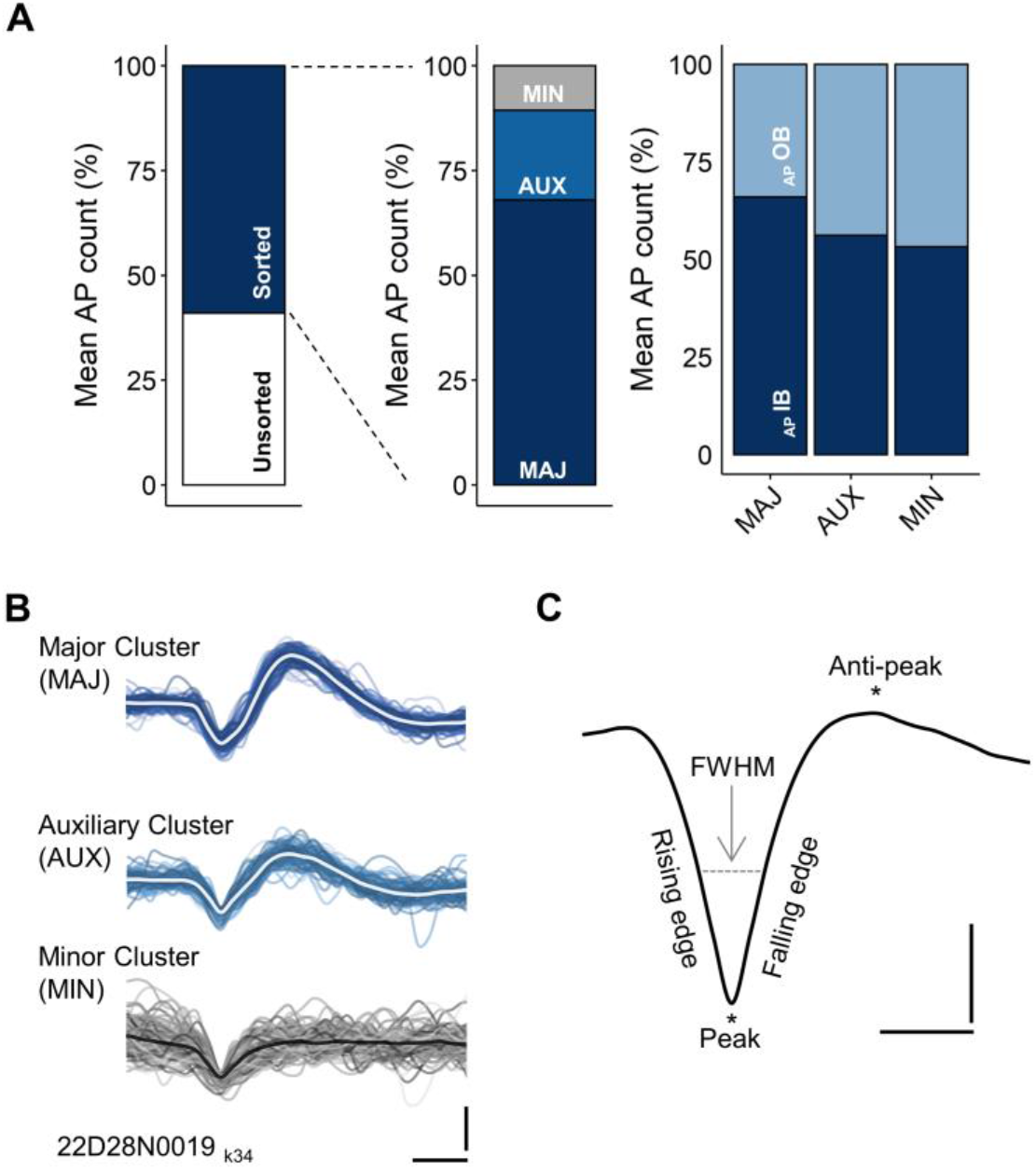
AP detection, sorting, cluster repartition and waveform analysis. **(A)** From left to right: Mean AP detection for unsorted and sorted AP fraction (% of total detected APs); Relative fraction of sorted APs attributed to Major (MAJ), Auxiliary (AUX) and Minors (MIN) clusters; Mean AP count for sorted AP occurring either inside (AP IB) or outside (AP OB) bursts period. Data collected over 15 min during the pre-exposure phase from 15 cultures of the RF group used here as representative. **(B)** Example of sorted AP waveforms after principal component analysis and hierarchical classification, overlay of 125 waveforms per cluster with averaged waveform highlighted, data from one channel of a the same culture. Scale: (y): 40 μV; (x): 500 μs. **(C)** Illustration of the metrics used to quantify changes in AP waveforms. FWHM: full width at half maximum. As recorded extracellularly the AP waveform is inverted. Scale: (y): 10 μV; (x): 500 μs.

**S3 Table.** Metrics definition: Analysis of AP waveform

| Metrics |  |
| --- | --- |
| Peak amplitude ($\mu\text{V}$) | Maximal amplitude of the first peak (negative going) |
| Anti-peak amplitude ($\mu\text{V}$) | Maximal amplitude of the second peak (positive going) |
| Half-width ($\mu\text{s}$) | Full width at half maximum (FWHM) of the first peak computed with linear interpolation |
| Rising edge ($\mu\text{V}/10^{-01}\text{ ms}$) | Maximal slope of the AP rising edge (negative going) |
| Falling edge ($\mu\text{V}/10^{-01}\text{ ms}$) | Maximal slope of the AP falling edge (positive going) |

**S4 Table.** Average raw values at baseline of the various metrics used to quantify change in AP waveform across the various experimental groups

| Metrics | SH | RF | MU |
| --- | --- | --- | --- |
| Channel per MEA (count) | 53 (12) | 54 (6.5) | 53 (2.3) |
| Sorted AP (count) | 23 266 (18 327) | 31 713 (17 353) | 139 499 (87 694) * |
| Peak amplitude ($\mu\text{V}$) | -27.3 (11.1) | -28.9 (6.2) | -29.1 (5.7) |
| Anti-peak amplitude ($\mu\text{V}$) | 11.2 (7.8) | 13.1 (6.5) | 15.3 (2) |
| Half-width ($\mu\text{s}$) | 224.6 (39.7) | 247.9 (29.8) | 253.5 (52.3) |
| Rising edge ($\mu\text{V}/10^{-01}\text{ ms}$) | -12.8 (5.7) | -13.7 (3) | -13.9 (4.8) |
| Falling edge ($\mu\text{V}/10^{-01}\text{ ms}$) | 12.8 (7.3) | 14.6 (3.8) | 15.21 (6) |

Data expressed as Median (IQR). \* Indicates significant difference ( $p < 0.05$ ) between MU-SH and MU-RF pairs. Pairwise comparison done with Kruskal-Wallis test followed by Conover's all-pairs posthoc test. SH, $n = 12$ ; RF, $n = 15$ ; MU, $n = 8$ .

